# Single-cell RNA sequencing reveals collagen interactions between detached osteoclasts and activated fibroblasts in granulation tissue surrounding sequestra in medication-related osteonecrosis of the jaw

**DOI:** 10.64898/2026.01.30.702745

**Authors:** Chika Nakanishi, Takeshi Yoshida, Takuo Emoto, Yasumasa Kakei, Daisuke Takeda, Yujiro Hiraoka, Manabu Shigeoka, Akira Kimoto, Takumi Hasegawa, Akishige Hokugo, Tetsuya Hara, Yoshiaki Tadokoro, Noriaki Emoto, Junichi Kikuta, Tomoya Yamashita, Masaya Akashi

## Abstract

**Objectives:** Medication-related osteonecrosis of the jaw (MRONJ) is a rare but serious complication of anti-resorptive agents used for osteoporosis and bone metastasis. As the global population ages, the number of patients with MRONJ continues to rise. Resorptive osteoclasts firmly attached to bone surfaces cannot be detected using single-cell RNA-Seq (scRNA-Seq); however, we hypothesized that multinucleated giant and detached osteoclasts (DetOCs) can be detected by this method. This study aimed to characterize DetOCs with scRNA-seq.

**Materials and Methods:** scRNA-Seq and immunohistochemistry staining was performed on granulation tissues surrounding sequestra from four patients with MRONJ and radicular cysts, a common odontogenic infectious disease with normal bone metabolism.

**Results:** scRNA-Seq and immunohistochemistry staining revealed that DetOCs were detected exclusively in the granulation tissues of MRONJ. These DetOCs expressed well-known osteoclast markers as well as *COL27A1*. Subcluster analysis revealed that *SPP1*^*+*^ *TREM2*^*+*^ macrophages differentiated into DetOCs and expressed several types of collagens. Immunostaining confirmed the expression of *COL27A1* not only in DetOCs in MRONJ but also in osteoclasts in mandibular cancer. CellChat analysis revealed interactions mediated by collagen signaling pathways between DetOCs, myeloid cells, and activated fibroblasts.

**Conclusions:** We successfully characterized DetOCs and identified a distinctive microenvironment involving DetOCs and activated fibroblasts in the granulation tissues surrounding sequestra in MRONJ.

**Clinical Relevance:** Identification of DetOCs and their unique interaction with activated fibroblasts in MRONJ lesions provides novel insight into the pathophysiology of sequestrum formation, suggesting potential therapeutic targets to ameliorate MRONJ.

## Introduction

Medication-related osteonecrosis of the jaw (MRONJ) is a serious adverse reaction to anti-resorptive agents (ARAs) such as bisphosphonates (BPs) and denosumab (an anti-receptor activator of nuclear factor kappa-B ligand [RANKL]) used in the treatment of osteoporosis and bone metastases in patients with cancer [1]. Although the incidence of MRONJ is considerably lower in patients with osteoporosis (<0.05%) than in those with cancer metastases (<0.5%) [2], the number of patients with osteoporosis who develop MRONJ cannot be overlooked—particularly in a super-aged society like Japan [3]. The diverse symptoms of MRONJ, including intractable pain, infection, insufficient oral intake, and pathological fracture, significantly impair patients’ quality of life [1]. Further elucidation of MRONJ pathology is still needed to develop more effective methods for its prevention and treatment.

In MRONJ, sclerosis with an unclear boundary is often observed around unhealed extraction sockets on X-ray or computed tomography (CT) images (Fig. 1a). As the condition progresses, osteolysis occurs in some cases, and the boundary between necrotic and vital areas may become clearer, eventually leading to sequestrum formation (Fig. 1a). Resection of sclerotic lesions with an unclear boundary often makes it difficult to determine the resection margin, resulting in uncertain surgical outcomes. By contrast, conservative surgery—such as sequestrectomy—tends to lead to better outcomes in cases where a sequestrum has formed [4]. Our previous study showed a statistically significant association between a poor prognosis in patients with MRONJ and the absence of sequestrum formation [5]. Thus, elucidating the underlying mechanisms of sequestrum formation remains important. In recent years, single-cell RNA-Seq (scRNA-Seq) has been used in various studies, but analyzing osteoclasts has been considered difficult because they are firmly attached to the bone surface. Notably, large numbers of multinucleated giant osteoclasts with prolonged lifespans affected by BPs have been observed in bone biopsy specimens from patients treated with BPs [6]. These abnormal osteoclasts do not adhere tightly to the bone surface, so we designate them as **<u>det</u>**ached **<u>o</u>**steo**<u>c</u>**lasts (DetOCs). We hypothesized that DetOCs might be involved in the granulation tissues surrounding the sequestrum and therefore conducted scRNA-Seq analysis. To clearly distinguish DetOCs in MRONJ from other cell types, we selected another common odontogenic infectious disease, the radicular cyst, as a control.Radicular cysts involve tartrate-resistant acid phosphatase (TRAP)-positive osteoclast precursors and a few runt-related transcription factor 2 (RUNX2)-positive osteoblastic cells [7,8]. Unlike MRONJ, in which osteoclast–osteoblast coupling is abnormal, normal bone coupling is maintained in radicular cysts. Therefore, we speculated that in radicular cysts, normal osteoclasts remain firmly attached to the bone surface, and thus—even if detectable by scRNA-Seq—only their precursor cells might be observed. This study aimed to characterize DetOCs with scRNA-seq. In fact, we successfully detected DetOCs, which were abundantly present only in MRONJ but not in radicular cysts. DetOCs highly express certain types of collagen. In MRONJ, activated fibroblasts increase in number and send collagen signals to DetOCs and myeloid cells, suggesting that corporation among these cells likely contributes to sequestrum formation.

**Fig. 1:**
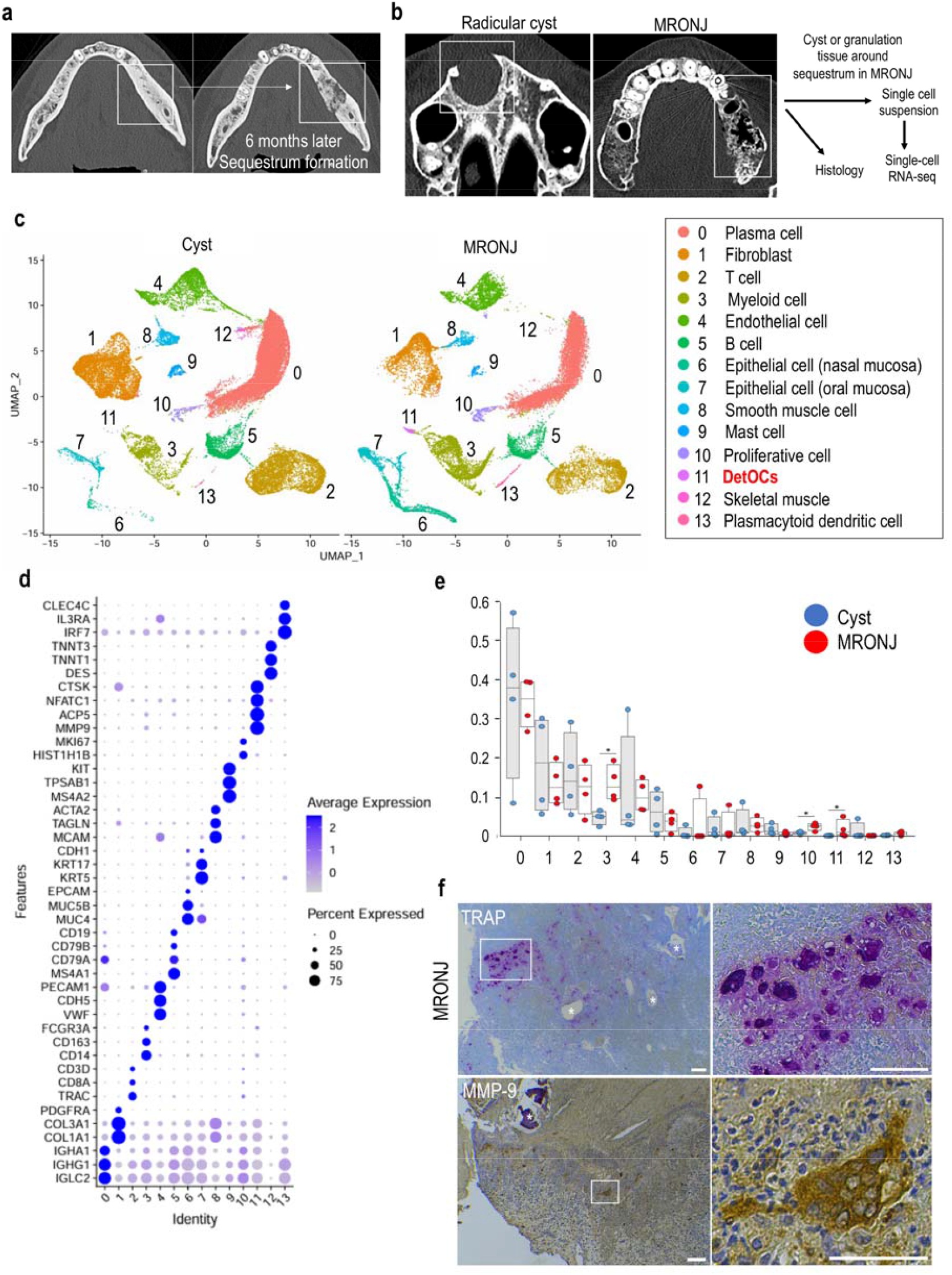
Detached osteoclasts were detected only in granulation tissues surrounding sequestra in MRONJ. **a** Representative CT images showing sequestrum formation in MRONJ. **b** Schematic illustration of the study design. Representative CT images are from a radicular cyst (Patient No. 2) and MRONJ (PatientNo. 1). **c** UMAP plot of scRNA-Seq transcriptome profiles from radicular cyst and MRONJ samples. **d** Dot plot showing the expression of selected marker genes for the cell subsets identified in **c. e** Proportions of each cell population in radicular cyst (blue) and MRONJ (red). Data are presented as median with interquartile range and were compared using the Wilcoxon rank-sum test. *p* < 0.01. **f** TRAP staining (upper panels) and MMP-9 staining (lower panels) of MRONJ tissues. White asterisks indicate sequestra. White boxes highlight enlarged views. Scale bar: 100 μm.

## Materials and Methods

### Patients and sample collection

Four patients who underwent surgery for MRONJ at our department between October and November 2023 were selected for scRNA-Seq. Granulation tissues surrounding the necrotic bone were collected as samples. As a control group, four patients who underwent surgery for radicular cysts between December 2023 and May 2024 were selected. The collected radicular cyst tissues were used as control samples.Additionally, six patients who underwent surgery for mandibular squamous cell carcinoma were randomly selected for immunostaining to evaluate COL27A1 expression.

The following epidemiologic data were gathered from patients’ medical charts: sex, age, lesion location (maxilla or mandible), underlying diseases, use of ARAs, comorbidities, and medication history. All patients provided written informed consent for the use of their clinical information and surgical samples, including granulation tissues around necrotic bone, for this study. These samples were used to prepare scRNA-Seq libraries.

### Histology

Tissue samples obtained during surgery were immediately washed in PBS and fixed in 4% paraformaldehyde, then processed using a standard paraffin-embedding protocol. Paraffin-embedded tissue sections were cut at a thickness of 4 μm and subjected to H&E staining and immunohistochemical staining. TRAP staining was performed using a TRAP/ALP Staining Kit (#294-67001; FUJIFILM Wako). After deparaffinization and rehydration, sections were incubated at 37°C for 30 minutes in TRAP staining solution composed of 1 mL tartaric acid solution, 9 mL substrate solution A, and 0.1 mL substrate solution B. Following rinsing in distilled water, nuclear staining was performed for 4–5 seconds, followed by additional washes. Slides were dried on a heating plate at 37°C, cleared with xylene, and coverslipped. Enzyme-based immunohistochemical staining was performed to detect DetOCs and myeloid cells. Antigen retrieval was conducted by autoclaving sections at 121°C for 10 minutes in either 1 mM Tris-EDTA buffer (pH 9.0) or 10 mM citrate buffer (pH 6.0), depending on the antigen. Endogenous peroxidase activity was blocked with 3% hydrogen peroxide for 15 minutes. After washing, sections were blocked with Blocking I solution and incubated overnight at 4°C with the respective primary antibodies: anti-MMP9 (rabbit polyclonal, clone RP066-01, 1:500; Diagnostic BioSystems), anti-DPP-4 (ab215711, rabbit monoclonal, clone EPR20819, 1:1000; Abcam),anti-osteopontin (ab166709, mouse monoclonal, clone 7C5H12, 1:100; Abcam), anti-CKB (#66764-1-Ig, mouse monoclonal, clone 2B7C3, 1:1000; Proteintech), anti-collagen type XXVII (#15673-1-AP, rabbit polyclonal, 1:50; Proteintech), anti-COL6A2 (#C12205, rabbit polyclonal, 1:200; Assay Biotechnology), anti-COL6A1 (#E9U3B, rabbit monoclonal, 1:100; Cell Signaling Technology), and anti-collagen V (#PA5-102416, rabbit polyclonal, 1:100; Invitrogen). Appropriate HRP-conjugated secondary antibodies were applied for 30 minutes at room temperature (anti-rabbit IgG: #424144, anti-mouse IgG: #424134;Nichirei). The DAB reaction was carried out for 1–5 minutes depending on the target, followed by Mayer’s hematoxylin counterstaining, dehydration, clearing, and mounting.

For double immunofluorescence staining targeting combinations of CKB/COL27A1, antigen retrieval was performed by autoclaving sections in either citrate buffer (pH□6.0) or Tris-EDTA buffer (pH□9.0). After blocking, sections were incubated overnight at 4°C with the same primary antibodies described above. Following washing, secondary antibodies conjugated to Alexa Fluor dyes were applied for 60 minutes at room temperature: Alexa Fluor 488 chicken anti-mouse IgG (A21200, 1:500; Invitrogen) and Alexa Fluor 594 chicken anti-rabbit IgG (A21442, 1:500; Invitrogen). Nuclei were counterstained with DAPI for 10 minutes at room temperature, and sections were mounted using aqueous mounting medium.

All immunostaining procedures were outsourced to Kyodo Byori (Kobe, Japan). The stained sections were examined using an all-in-one fluorescence microscope (BZ-X700; Keyence).

### Sample preparation for single-cell analysis

Fresh granulation tissue samples were collected at the time of surgery. The tissue was washed with PBS, dehydrated, and immediately flash-frozen in liquid nitrogen. Bone fragments mixed within the granulation tissue were carefully removed. The frozen tissue samples were stored in a liquid nitrogen tank and shipped in a shipping container to a contract laboratory. Tissue dissociation and fixation were performed at the contract laboratory according to the Chromium Fixed RNA Profiling workflow (CG000553 Rev. B; 10x Genomics). Dissociation was carried out using a gentleMACS Octo Dissociator with Heaters (Miltenyi Biotec), and single-cell suspensions were prepared through combined mechanical and enzymatic treatment. The resulting cells were fixed with formaldehyde.

### Single-cell sequencing library preparation

Single-cell gene expression profiling was performed using the Chromium Fixed RNA Profiling workflow (CG000527; 10x Genomics). Library preparation involved the Chromium Fixed RNA Kit (Human Transcriptome) and the Chromium Next GEM Single Cell Fixed RNA Sample Preparation Kit. Following hybridization of detection probes to RNA within fixed cells, adjacent probes were joined through a ligation reaction. Samples were then pooled to normalize cell counts across groups and processed on the Chromium X instrument, where cells were encapsulated with barcoded primer beads. Annealing and extension reactions added cell-identifying adapter sequences, and indexed primers were used for PCR amplification and library purification, generating sequencing-ready libraries.

### ScRNA-Seq data processing

The prepared libraries were pooled and sequenced on an Illumina NovaSeq X Plus using 150 bp paired-end reads, generating approximately 2 billion reads (approximately 300 Gb total). Demultiplexing was performed using Cell Ranger mkfastq (v7.2.0) and bcl2fastq2 (v2.20), producing FASTQ files that underwent quality assessment. Raw sequencing data were processed into expression matrices using default parameters in Cell Ranger single-cell software (v7.2.0; 10x Genomics) with the GRCh38 human reference genome. Prior to downstream analysis, the resulting count matrices were processed with CellBender remove-background (v0.3.0) to identify non-empty droplets and remove ambient RNA using customized parameters [9]. Data analysis was carried out using Seurat (v4.1.1) within the R 4.2.1 environment [10]. Mitochondrial genes were removed during initial filtering, and only cells expressing between 300 and 6000 genes, with mitochondrial gene content below 6%, and expressing each gene in at least three cells were retained to exclude doublets and low-quality cells.

Data integration and batch effect correction were performed using the reciprocal principal component analysis (“rpca”) method within Seurat’s integration workflow, followed by log-normalization with default settings. Clustering was performed on the integrated gene expression matrix using Seurat with a resolution parameter of 0.1. For dimensionality reduction, principal component analysis was performed, and the first 20 principal components were used for uniform manifold approximation and projection (UMAP) with default parameters [11]. Differential gene expression was assessed using the nonparametric Wilcoxon rank-sum test implemented in JMP version 13.1.0 (SAS Institute Inc., Cary, NC, USA). By comparing gene expression in each cluster to that of all other clusters, the top 10 differentially expressed genes unique to each cluster were identified and visualized as heatmaps. Dot plots were also used to display the expression of cluster-specific marker genes or known standard markers. Feature plots were used to visualize the expression patterns of selected genes on a UMAP embedding, allowing assessment of the spatial distribution of gene expression across cell clusters.

We subsequently extracted two clusters of interest, namely myeloid cells and DetOCs, from the integrated dataset. After merging these two clusters, re-clustering was performed using a resolution parameter of 0.34, initially generating 13 clusters. To reduce redundancy and simplify interpretation, clusters with similar gene expression profiles were merged: clusters 1 and 3, clusters 2 and 10, and clusters 5, 6, and 11 were combined, resulting in a total of nine distinct clusters. Re-clustering was similarly performed for fibroblasts. For fibroblasts, subclustering was conducted with a resolution parameter of 0.1, generating seven clusters. Cluster 5 was excluded from further analysis because of suspected T-cell doublets, evidenced by high expression of T-cell markers such as *TRAC* and *TRBC1*. After removal of this cluster, six fibroblast clusters remained for downstream analysis.

### RNA velocity analyses

In our dataset, splicing information could not be obtained because of the fixation process used during sample preparation. Therefore, RNA velocity analysis was performed using TFvelo, a method that extends the concept of RNA velocity to various single-cell datasets without relying on splicing information by incorporating gene regulatory information. Using SeuratDisk, the single-cell data prepared in Seurat were converted into the Python data format, anndata, enabling visualization of the estimated velocities on UMAP plots generated from the Seurat analysis [12].

### Pseudotime gene expression analysis

Gene expression of *COL27A1* was plotted along pseudotime using a smoothed curve generated by CellRank2, with each dot representing a single cell.

### Gene ontology analyses

GO analysis for biological processes was performed using the clusterProfiler package, based on the differentially expressed genes identified in each cluster of myeloid cells [13].

### CellChat analyses

Cell–cell interaction analysis was performed using the CellChat package (version 1.6.0), which was used to infer and evaluate the signaling pathway network of clusters fibroblasts and myeloid cells [14].

### Statistics and reproducibility

The Wilcoxon rank-sum test was used to compare gene expression levels between clusters or sub-clusters. All statistical tests were two-sided, and *p* < 0.05 was considered statistically significant.

### Study approval

Eight patients with radicular cysts and MRONJ were prospectively recruited for this study. Written informed consent was obtained from all participants. The study was approved by the Ethics Committees of Kobe University (B230094) and conformed to the principles of the Declaration of Helsinki.

### Data availability

All sequencing data are publicly available at GSE303003.

## Results

### Detached osteoclasts only in the granulation tissues surrounding sequestra in MRONJ

To perform transcriptome profiling of human radicular cysts and granulation tissues in MRONJ, surgical specimens were obtained under general anesthesia. The subsequent sample processing is described in the Methods section.

The patients’ characteristics and preoperative CT images are shown in Table 1 and Supplemental Fig. 1a. As shown in Fig. 1b, the bone lining around the cyst was preserved in radicular cysts because coupling between bone resorption and formation remained unaffected. By contrast, bone resorption areas around the sequestrum in MRONJ exhibited an unclear boundary, lacking the bone lining seen in radicular cysts, secondary to the effects of BPs. Sequestrum formation was detected in the preoperative CT images of all patients (Supplemental Fig. 1a).

**Table 1.**
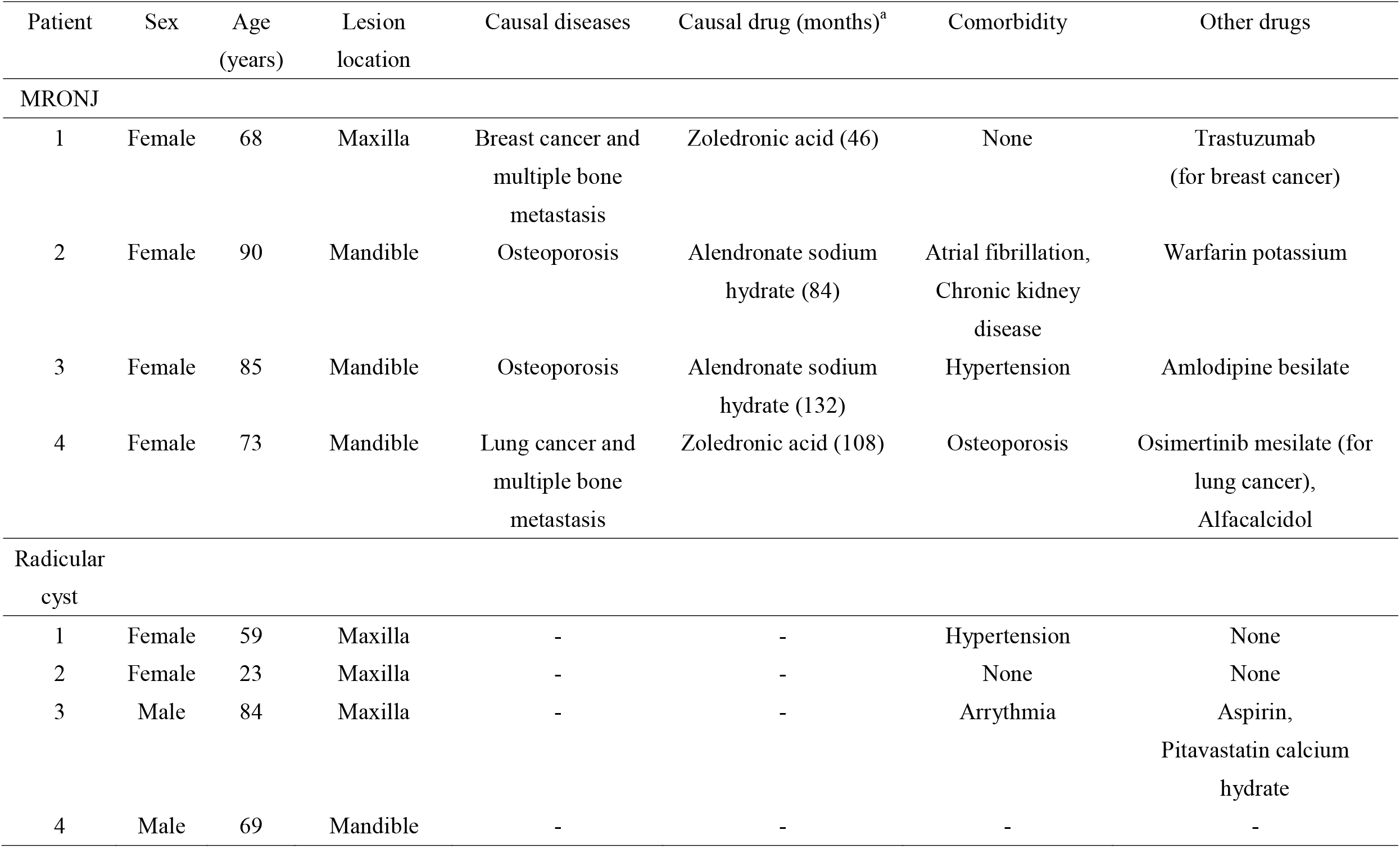
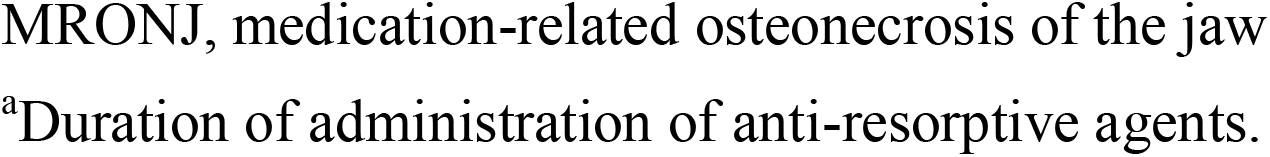
Characteristics of patients with MRONJ and radicular cysts.

| Patient | Sex | Age<br>(years) | Lesion<br>location | Causal diseases | Causal drug (months)<sup>a</sup> | Comorbidity | Other drugs |
| --- | --- | --- | --- | --- | --- | --- | --- |
| MRONJ |  |  |  |  |  |  |  |
| 1 | Female | 68 | Maxilla | Breast cancer and multiple bone metastasis | Zoledronic acid (46) | None | Trastuzumab (for breast cancer) |
| 2 | Female | 90 | Mandible | Osteoporosis | Alendronate sodium hydrate (84) | Atrial fibrillation, Chronic kidney disease | Warfarin potassium |
| 3 | Female | 85 | Mandible | Osteoporosis | Alendronate sodium hydrate (132) | Hypertension | Amlodipine besilate |
| 4 | Female | 73 | Mandible | Lung cancer and multiple bone metastasis | Zoledronic acid (108) | Osteoporosis | Osimertinib mesilate (for lung cancer),<br>Alfacalcidol |
| Radicular cyst |  |  |  |  |  |  |  |
| 1 | Female | 59 | Maxilla | - | - | Hypertension | None |
| 2 | Female | 23 | Maxilla | - | - | None | None |
| 3 | Male | 84 | Maxilla | - | - | Arrhythmia | Aspirin,<br>Pitavastatin calcium hydrate |
| 4 | Male | 69 | Mandible | - | - | - | - |

MRONJ, medication-related osteonecrosis of the jaw
<sup>a</sup>Duration of administration of anti-resorptive agents.

After standard data processing and quality filtering, single-cell transcriptomes from a total of 60041 cells—33309 from radicular cysts and 26732 from the granulation tissues in MRONJ—were obtained. We identified 14 clusters based on transcriptional similarity, which we labeled according to the expression of well-known marker genes (Fig. 1c, d). The structural cells included {4} blood endothelial cells (characterized by *PECAM1, CDH5*, and *VWF*); {6} epithelial cells of the nasal mucosa (*MUC4* and *MUC5B*), found only in maxillary cases; {7} epithelial cells of the oral mucosa (*KRT5* and *KRT17*) [15];{8} smooth muscle cells (*MCAM* and *ACTA2*) [16]; and {12} skeletal muscle cells (*DES* and *TNNI1*) [17,18]. {1} Fibroblasts were characterized by *COL1A1* expression, and {10} proliferative cells by *MKI67* [19]. Among the immune cells, we distinguished {0} plasma cells (*IGHG1*) [19], {2} T cells (*TRAC* and *CD8A*) [19], {3} myeloid cells (*CD14* and *CD163*) [20], {5} B cells (*MS4A1* and *CD79A*) [21], {9} mast cells (*TPSAB1*) [19,20], and {13} plasmacytoid DCs (*CLEC4C*) [22]. Cluster {11} consisted of cells characterized by expression of major osteoclastic markers such as *CTSK, NFATC1*, and *ACP5*, and these were designated DetOCs [19]. The proportions of cluster {11} DetOCs (*p* = 0.02),{3} myeloid cells (*p* = 0.02), and {10} proliferative cells (*p* = 0.04) were significantly higher in MRONJ than in radicular cysts (Fig. 1e). We confirmed the presence of multinucleated giant TRAP- and MMP-9-positive cells within the granulation tissues in MRONJ and observed that these cells did not adhere to the bone surface but appeared to float around the sequestra (Fig. 1f).

### Characterization of detached osteoclasts

Next, we performed subclustering of myeloid cells ({3} in total cells) and DetOCs ({11}) to provide a more detailed characterization of DetOCs. We identified nine clusters based on transcriptional similarity using automated cell-type annotation (Fig. 2a and Supplemental Fig. 2a): {My 0} SELENOP^+^ resident macrophages characterized by expression of *SELENOP* and *C1QA* [23]; {My 1} classical monocytes characterized by expression of *IL1*β and *S100A9* [24,25]; {My 2} CCL18^+^ macrophages, known as an inflammation-suppressive subgroup [26]; {My 3} SPP1^+^ TREM2^+^ macrophages, associated with pro-fibrotic potential [27]; {My 4} neutrophils characterized by expression of *G0S2* [28]; {My 5} DetOCs; {My 6} conventional DCs characterized by expression of *CLEC9A* and *XCR1* [29]; {My 7} mature DCs enriched in immunoregulatory molecules characterized by expression of *CD40* and *CCR7* [30]; and {My 8} proliferative cells. A radicular cyst is an inflammatory and bone-resorptive disease in which bone coupling is not affected by ARAs. Therefore, neither DetOCs nor normal osteoclasts, which were likely tightly attached to the bone surface, were detected in radicular cysts.

**Fig. 2:**
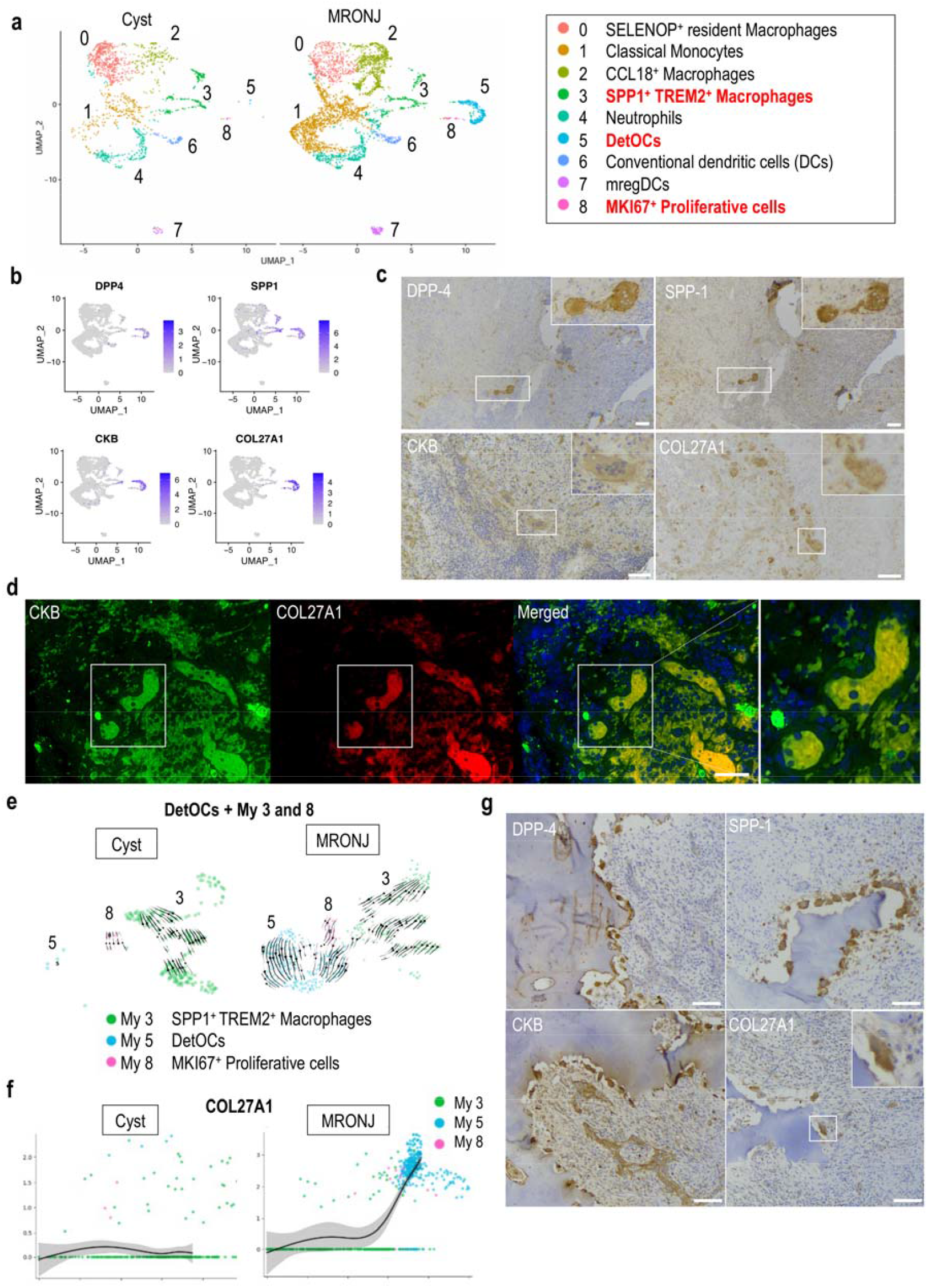
Characterization of detached osteoclasts. **a** UMAP of scRNA-Seq transcriptome profiles from myeloid cells ({3} in Fig. 1c) and DetOCs ({11} in Fig. 1c) from radicular cyst and MRONJ samples. **b** Feature plots showing *DPP4, SPP1, CKB*, and *COL27A1* expression in myeloid cells and DetOCs. **C** Immunostaining for DPP-4, SPP-1, CKB, and COL27A1 in MRONJ tissues. Enlarged views show giant multinucleated cells, likely DetOCs. Scale bar: 100 μm. **d** Double immunofluorescence staining for CKB and COL27A1 in MRONJ specimens. Enlarged views highlight CKB- and COL27A1-positive DetOCs. Scale bar: 50 μm. **e** RNA velocity field projected onto UMAP plots of {My 3} SPP1^+^ TREM2^+^ macrophages, {My 5} DetOCs, and {My 8} MKI67^+^ proliferative cells in radicular cysts and MRONJ. **f** RNA velocity analysis demonstrating upregulation of *COL27A1* in DetOCs specifically within MRONJ. **g** Immunostaining for DPP-4, SPP-1, CKB, and COL27A1 in mandibular cancer tissue. Enlarged views show osteoclasts. Scale bar: 100 μm.

The proportions of {My 5} DetOCs and {My 2} CCL18^+^ inflammation-suppressive macrophages were significantly higher in MRONJ than in radicular cysts, whereas the proportion of {My 0} SELENOP^+^ resident macrophages was higher in radicular cysts (Supplemental Fig. 2b). DetOCs highly expressed well-known osteoclast markers such as *ACP5, CTSK*, and *NFATC1* (Supplemental Fig. 2a–c). The heatmap (Supplemental Fig. 2a) and feature plots (Fig. 2b) revealed that *DPP4, SPP1, CKB*, and *COL27A1* were also highly expressed in DetOCs. While *DPP4* has been suggested to play a role in osteoclast–osteoblast coupling [31] and *CKB* has been reported to be involved in osteoclast-mediated bone resorption [32], no previous reports have described the role of *COL27A1* in osteoclasts. The expression of *COL27A1* was found not only in DetOCs but also in fibroblasts, epithelial cells, and smooth muscle cells in the total cell population, as described below. We confirmed the expression of DPP-4, SPP-1, CKB, and COL27A1 proteins in multinucleated giant cells around necrotic bone using immunostaining of surgical specimens (Fig. 2c). Double immunofluorescence staining further revealed the co-expression of COL27A1 and CKB proteins in the multinucleated giant cells (Fig. 2d).

Next, we examined COL27A1 expression in DetOCs. We investigated the differentiation into DetOCs, with a particular focus on collagen expression. RNA velocity was analyzed by distinguishing between unspliced and spliced mRNAs [33]. In cysts, the differentiation direction of both {My 3} SPP1^+^ TREM2^+^ macrophages and {My 8} MKI67^+^ proliferative cells pointed toward an undetected cell population, whereas in MRONJ, {My 3} and {My 8} differentiated into {My 5} DetOCs (Fig. 2e). Pseudotime gene expression analysis focusing on *COL27A1* revealed that its expression was markedly upregulated in DetOCs in MRONJ (Fig. 2f). Moreover, we investigated whether *COL27A1* expression is specific to DetOCs in the unique environment of MRONJ or whether it also appears in other conditions. We evaluated osteoclasts in mandibular gingival squamous cell carcinoma and found that COL27A1 was also expressed in active bone-resorbing osteoclasts in cancer, along with DPP-4, SPP-1, and CKB (Fig. 2g). Importantly, the DPP-4-, SPP-1-, CKB-, and COL27A1-positive cells observed in mandibular cancer exhibited morphology resembling that of normal osteoclasts, in contrast to the multinucleated giant DetOCs seen in MRONJ.

### Characteristic expression of collagens in detached osteoclasts, myeloid cells, and fibroblast in MRONJ

Unexpectedly, we found *COL27A1* expression in DetOCs and therefore evaluated collagen signaling in DetOCs and myeloid cells. Gene ontology (GO) enrichment analysis revealed that macrophage-, osteoclast-, and collagen-related pathways differed between cysts and MRONJ (Fig. 3a). Previous reports have suggested that macrophages express collagen and that the type of collagen expressed may reflect macrophage characteristics [34-36]. We comprehensively investigated the expression of all types of collagen in DetOCs and myeloid cells and found that {My 5} DetOCs expressed not only *COL27A1* but also *COL6A2, COL6A1*, and *COL5A1*, while {My 3} SPP1^+^ TREM2^+^ macrophages expressed *COL3A1, COL1A2*, and *COL1A1* (Fig. 3b).

**Fig. 3:**
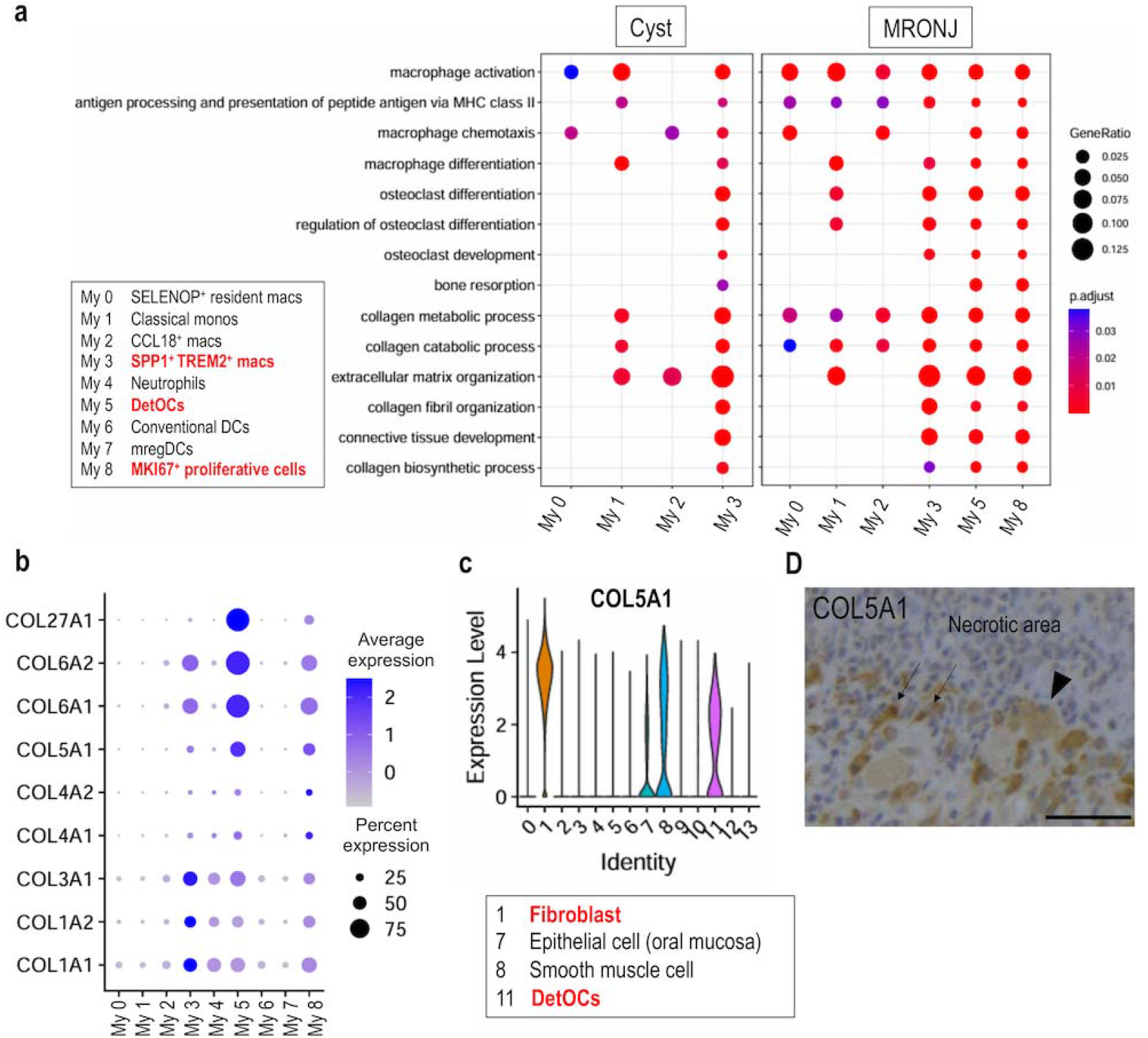
Characteristic expression of collagens in detached osteoclasts, myeloid cells, and fibroblasts in MRONJ. **a** GO terms highlighting differences in macrophage- and osteoclast-related pathways between DetOCs and myeloid cells. **b** Dot plot showing collagen expression in DetOCs and myeloid cells. **c** Violin plot showing *COL5A1* expression in total cells. **d** Immunostaining for COL5A1 in MRONJ. Black arrowhead indicates COL5A1-positive multinucleated giant cells, and black arrows indicate COL5A1-positive spindle-shaped cells. Scale bar: 50 μm.

Violin plots of the total cell population reveled that *Col6A1* and *COL6A2* were expressed in DetOCs {11}, smooth muscle cells {8}, endothelial cells {4}, epithelial cells {6, 7} and fibroblasts {1}, whereas *Col5A1* was expressed in DetOCs {11}, smooth muscle cells {8}, and fibroblasts {1} (Supplemental Fig. 3a–c). We confirmed the presence of multinucleated giant COL5A1-positive cells at the boundary between necrotic areas and granulation tissues in MRONJ, with COL5A1-positive cells exhibiting spindle morphology near the multinucleated giant cells (Fig. 3d), as well as abundant COL6A1- and COL6A2-positive endothelial cells (Supplemental Fig. 3b). These findings indicate the accumulation of fibroblasts around DetOCs and myeloid cells around the sequestra.

### Collagen signals between detached osteoclasts, myeloid cells, and fibroblasts in MRONJ

Recent research has reported interactions between SPP1^+^ macrophages and FAP^+^ fibroblasts in cancer [37]. We were interested in the interaction between myeloid cells, including DetOCs, and fibroblasts; therefore, we performed subclustering of fibroblasts ({1} in total cells). Six clusters were identified based on transcriptional similarity using automated cell-type annotation (Fig. 4a): {Fb 0} RUNX2^+^ COL11A1^+^ mesenchymal fibroblasts [38], {Fb 1} FAP^+^ activated fibroblasts [37], {Fb 2} SPARC^+^ COL3A1^+^ fibroblasts known to have vascular interaction potential [39], {Fb 3} EGR1^+^ fibroblasts [40], {Fb 4} SFRP2^+^ COL15A1^+^ myofibroblast progenitors [41,42], and {Fb 5} COL18A1^+^ CD9^+^ fibroblasts [38,43] (Supplemental Fig. 4a–b). The proportion of {Fb 1} FAP^+^ activated fibroblasts was significantly higher in MRONJ than in radicular cysts, while the proportion of {Fb 3} EGR1^+^ fibroblasts was higher in radicular cysts (Supplemental Fig. 4c). Under normal conditions, *TNFSF11* (RANKL) signaling regulates bone metabolism [44]. We found that {Fb 0} mesenchymal fibroblasts highly expressed *RUNX2* but not *SP7*, indicating that {Fb 0} does not represent the typical RUNX2^+^ SP7^+^ preosteoblasts [45]. Furthermore, we found that DetOCs and some myeloid cell clusters ({My 7, 8}) expressed *TNFRSF11A* (RANK), but none of the fibroblast clusters expressed *TNFSF11* (RANKL), suggesting that RANKL signaling is not associated with DetOC formation. RNA velocity analysis revealed that fibroblast differentiation directions were scattered in cysts, while in MRONJ, all fibroblast populations ({Fb 0, 2, 3, 4}) converged into {Fb 1} FAP^+^ activated fibroblasts (Fig. 4b).

**Fig. 4:**
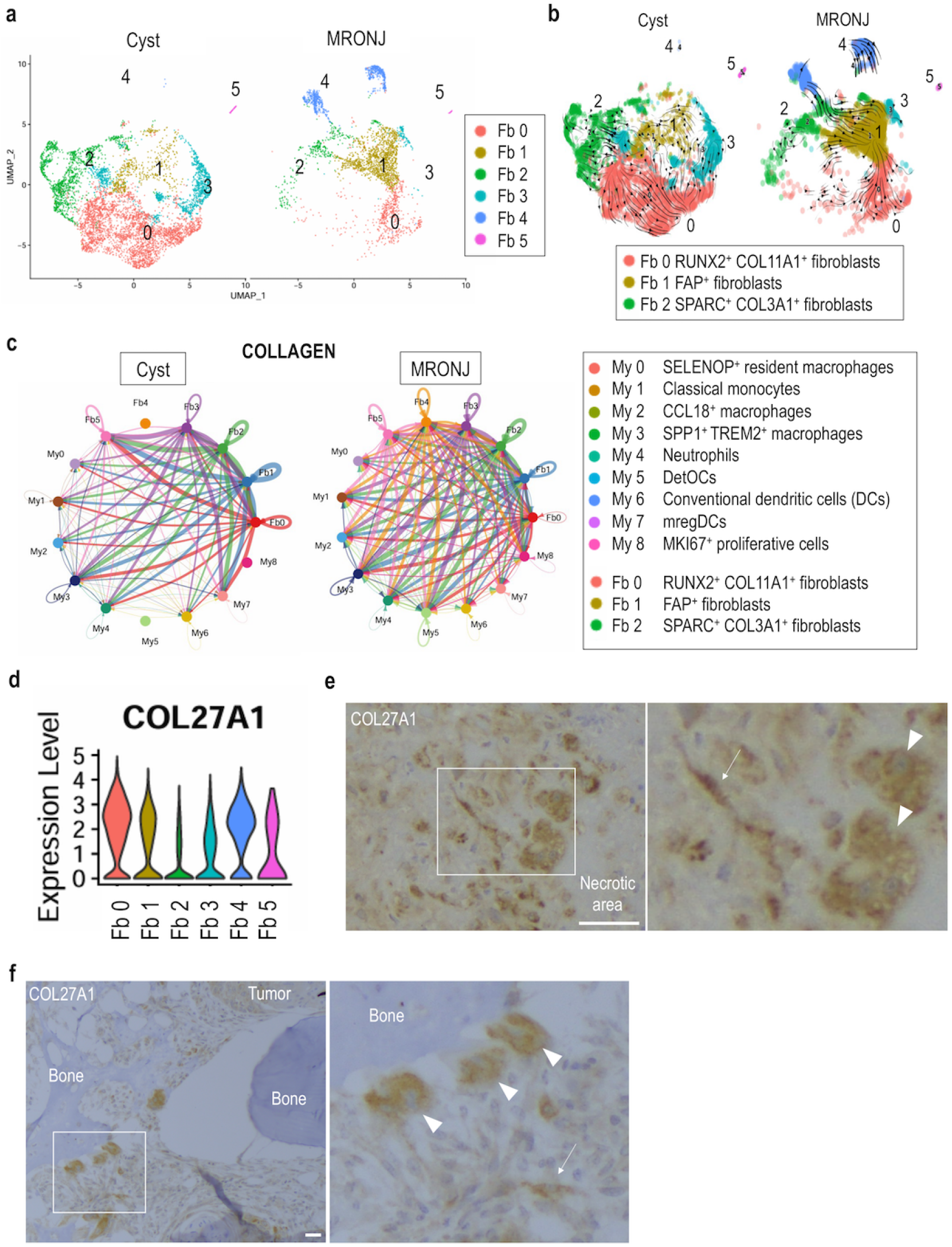
Collagen signals between detached osteoclasts, myeloid cells, and fibroblasts in MRONJ. **a** UMAP of scRNA-Seq transcriptome profiles from fibroblasts ({1} in Fig. 1c). **b** RNA velocity field projected onto UMAP plots of fibroblasts in radicular cysts and MRONJ. **c** CellChat analysis showing collagen signaling interactions between DetOCs, myeloid cells, and fibroblasts. **d** Violin plot showing *COL27A1* expression in fibroblast clusters. **e** Immunostaining for COL27A1 in MRONJ specimens. Enlarged view shows COL27A1-positive multinucleated giant cells (white arrowheads) and a spindle-shaped cell (arrow). Scale bar: 100 μm. **f** Immunostaining for COL27A1 in mandibular cancer tissue. Enlarged view shows COL27A1-positive osteoclasts (white arrowheads) and a spindle-shaped cell (arrow). Scale bar: 100 μm.

Finally, we sought to identify signaling pathways between DetOCs and myeloid cells using CellChat analysis. Interestingly, {Fb 1} FAP^+^ activated fibroblasts sent collagen signals mainly to {My 5} DetOCs and {My 3} SPP1^+^ TREM2^+^ macrophages in MRONJ (Fig. 4c). The violin plot revealed that COL27A1 was highly expressed in fibroblasts (Fig. 4d). We found COL27A1-positive multinucleated giant cells and spindle-shaped cells at the boundary between necrotic areas and granulation tissue areas (Fig. 4e). We also observed an accumulation of COL27A1-positive multinucleated giant cells and spindle-shaped cells at sites of active bone resorption in mandibular cancer (Fig. 4f).

## Discussion

MRONJ is a rare but serious complication of ARA treatment for osteoporosis and bone metastasis. As the global population ages, the growing number of patients affected by MRONJ should not be overlooked. Active osteoclasts have traditionally been considered difficult to analyze using scRNA-Seq because of their firm adhesion to bone surfaces. However, it is known that the multinucleated giant cells seen in osteonecrosis of the jaw are not firmly attached to bone surfaces because of the effects of ARAs and instead float near these surfaces. We focused on this characteristic and performed scRNA-Seq on granulation tissue surrounding the sequestrum in MRONJ, successfully detecting abnormal osteoclasts that were not adherent to bone surfaces, which we designated DetOCs. DetOCs not only highly expressed well-known osteoclast markers but also showed elevated expression of *COL27A1*, which has not been previously reported. In MRONJ, activated fibroblasts increased in number and sent collagen signals to DetOCs and myeloid cells.

It was previously believed that macrophages do not produce collagens, the major structural proteins of the intact extracellular matrix, but Schnoor et al.[34] showed that monocytes and macrophages express almost all known collagen and collagen-related mRNAs and that TGF-β1 strongly induces type VI collagen synthesis in vitro. Piccinini et al. [35] investigated the effects of distinct microenvironmental cues on macrophages differentiated from primary human monocytes and found that 24-hour stimulation with Gram-negative bacterial LPS and the C-terminal fibrinogen-like globe domain of tenascin-C upregulated the expression of various collagen mRNAs, including *COL27A*1. More recently, Sárvári et al. [36] performed single-nucleus RNA-Seq on epididymal white adipose tissue from mice fed either a low-fat or high-fat diet and identified six macrophage subpopulations: perivascular-like macrophages, lipid-associated macrophages, non-perivascular-like macrophages, proliferating lipid-associated macrophages, regulatory macrophages, and collagen-expressing macrophages. The authors speculated that collagen-expressing macrophages may play a role in extracellular matrix deposition and tissue remodeling due to the expression of collagen genes such as *Col3a1* and *Col5a2* [36]. In our study, DetOCs highly expressed *COL5A1* but not *COL5A2*, while SPP1^+^ TREM2^+^ macrophages highly expressed *COL3A1*. The function of collagen-expressing macrophages remains unclear, and to our knowledge, no previous studies have reported collagen expression in osteoclasts. Pham et al. [46] investigated *Col6a2*-knockout mice and found no differences in trabecular bone formation compared with wild-type mice; however, they observed an increased abundance of TRAP-positive osteoclasts in femoral sections of *Col6a2*-knockout mice, indicating that the primary effect of *COL6A2* deficiency is on osteoclastogenesis. The authors proposed an inhibitory function of *COL6A2*, suggesting that collagen VI produced by bone marrow stromal cells binds extracellular TNF-α and reduces its ability to promote osteoclastogenesis [46]. By contrast, DetOCs in our study themselves highly expressed *COL6A2*. Further investigations focusing on collagen expression in osteoclasts are warranted.

We found that DetOCs expressed collagen genes, among which *COL27A1* was the most highly expressed. To our knowledge, no previous studies have reported the expression of *COL27A1* in osteoclasts; however, one earlier study using murine bone marrow-derived macrophages showed that *COL27A1* mRNA transcription was induced by both LPS and heat-killed *Candida albicans* [47].*COL27A1* belongs to the fibrillar collagen gene family and is typically expressed in cartilaginous tissues [48]. The primary deposition site of the collagen encoded by *COL27A1* in human fetal skeletal tissue is at the transition zones from cartilage to bone [48]. Mutations in *COL27A1* are known to cause Steel syndrome, characterized by distinct facial features, short stature, irreducible bilateral hip and radial head dislocation, and carpal bone coalition [49]. Interestingly, a GWAS identified significant associations between SNPs in *GDF5* and *COL27A1* and the risk of knee pain [50]. Another recent study using the UK Biobank reported that *COL27A1* is among the candidate genes linked to undiagnosed diseases involving seizures and that compound heterozygous variants in *COL27A1* may underlie early-onset skeletal and skull abnormalities in these patients [51]. Unexpectedly, our immunostaining revealed that not only DetOCs in MRONJ but also osteoclasts in cancer tissues express COL27A1 protein. It has been considered difficult to analyze active osteoclasts using scRNA-Seq because of their rigid adhesion to the bone surface during resorption. However, several previous studies using scRNA-Seq in osteosarcoma have successfully identified osteoclasts [52,53]. Sun et al. [53] identified six distinct subgroups of osteoclasts in human osteosarcoma, including an *MKI67*^*+*^ osteoclast population. Because osteoclasts in cancer tissues appear to adhere less firmly to bone surfaces than do osteoclasts under normal physiological conditions, this may have enabled scRNA-Seq analysis similar to that used for MRONJ. Osteoclasts within cancer tissues exhibit this weak bone adherence, resembling the behavior of DetOCs in MRONJ. Further investigations are needed to clarify the role of *COL27A1* in DetOCs in MRONJ as well as in osteoclasts within cancerous tissues.

The primary objective of this study was to elucidate the underlying mechanism of sequestrum formation and progression in MRONJ. We found that no cell populations expressed RANKL, whereas DetOCs and myeloid cells expressed RANK, indicating that granulation tissues around sequestra in MRONJ lack RANKL signaling. Therefore, we sought to identify the signaling pathways involved in this RANKL-deficient environment. We found COL6A1-, COL6A2-, and COL5A1-positive multinucleated giant cells as well as an accumulation of COL6A1-, COL6A2-, and COL5A1-positive spindle-shaped cells near the DetOCs. Because we were interested in the role of spindle-shaped cells, we examined their characteristics and found that FAP^+^ activated fibroblasts were increased in MRONJ. CellChat analysis revealed that FAP^+^ activated fibroblasts mainly sent collagen signals to {My 5} DetOCs and {My 3} SPP1^+^ TREM2^+^ macrophages in MRONJ (Fig. 4c). In tumorigenesis, collagen produced by cancer-associated fibroblasts, including type XXVII collagen, has attracted increasing attention [54]. In our study, COL27A1 expression was detected in fibroblasts in the tumor microenvironment (Fig. 4f) and in spindle-shaped cells near COL27A1-positive multinucleated giant cells in MRONJ (Fig. 4e). These findings suggest that evaluating COL27A1 expression throughout the entire tissue, including diverse interstitial cells rather than focusing on specific cell types, may contribute to understanding and assessing the pathophysiology of MRONJ and sequestrum formation.

Finally, we must note the limitations of our study. First, the study was limited by a small sample size, and patients treated with denosumab could not be included. Second, we focused only on myeloid cells and fibroblasts, and the roles of other cell types, such as B cells, T cells, and epithelial cells, remain to be explored in future studies. Third, the findings presented here have not yet been validated through experiments using animal models or cultured cells. Fourth, the current data alone provide limited clinical implications. Recent preclinical studies have identified new osteoclast lineage cells, so-called osteomorphs, and dynamic osteoclcast recycling via osteomorphs [55]. However, this interesting osteoclast lineage cell has been identified only in mice [56]. Importantly, Park-Min et al. [56] indicated that the non-bone surface-associated TRAP-positive multinucleated cells observed following pamidronate treatment in patients [6] may interact with osteomorphs, an issue requiring detailed preclinical investigation. We believe that these “quiescent” TRAP-positive multinucleated cells described by Park-Min et al. [56] are indeed DetOCs themselves. Thus, our findings may help promote investigations into osteoclast recycling and the characterization of osteomorphs in humans.

In conclusion, we successfully identified an intriguing cell population—so-called DetOCs—in the granulation tissue surrounding sequestra in MRONJ. DetOCs exhibited high expression of well-known osteoclast markers and also showed elevated expression of *COL27A1*. Interestingly, immunohistochemical analysis revealed that COL27A1 is also expressed in osteoclasts within cancer tissues, suggesting the possibility that osteoclasts could be subclustered based on the types of collagen they express. In MRONJ, activated fibroblasts increased in number and sent collagen signals to DetOCs and myeloid cells. A key task for future research will be to elucidate how myeloid cells, including DetOCs, and activated fibroblasts contribute to the onset and progression of sequestrum formation in MRONJ.

## Acknowledgment

We thank Angela Morben, DVM, ELS, from Edanz (https://jp.edanz.com/ac), for editing a draft of this manuscript.

