## Supplementary figures and images for "Single-cell RNA sequencing reveals collagen interactions between detached osteoclasts and activated fibroblasts in granulation tissue surrounding sequestra in medication-related osteonecrosis of the jaw"

### Supplemental Figure 1

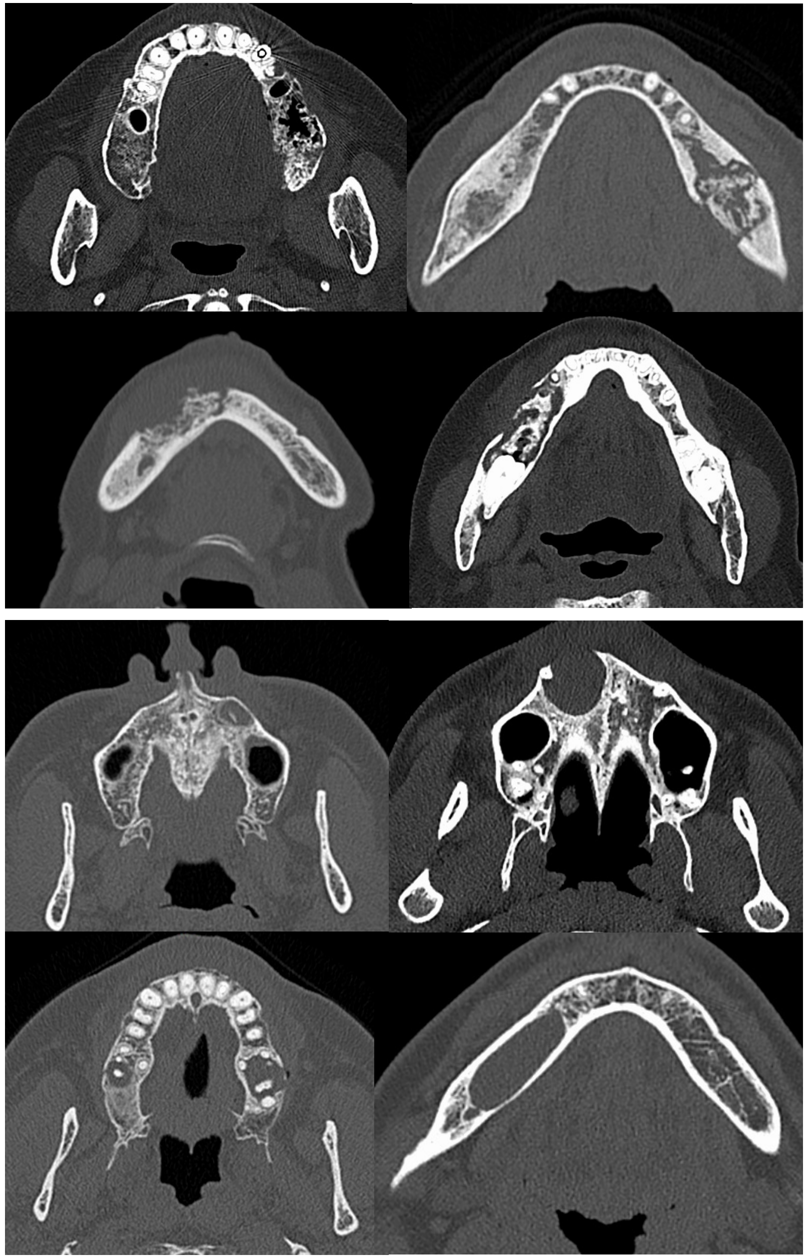

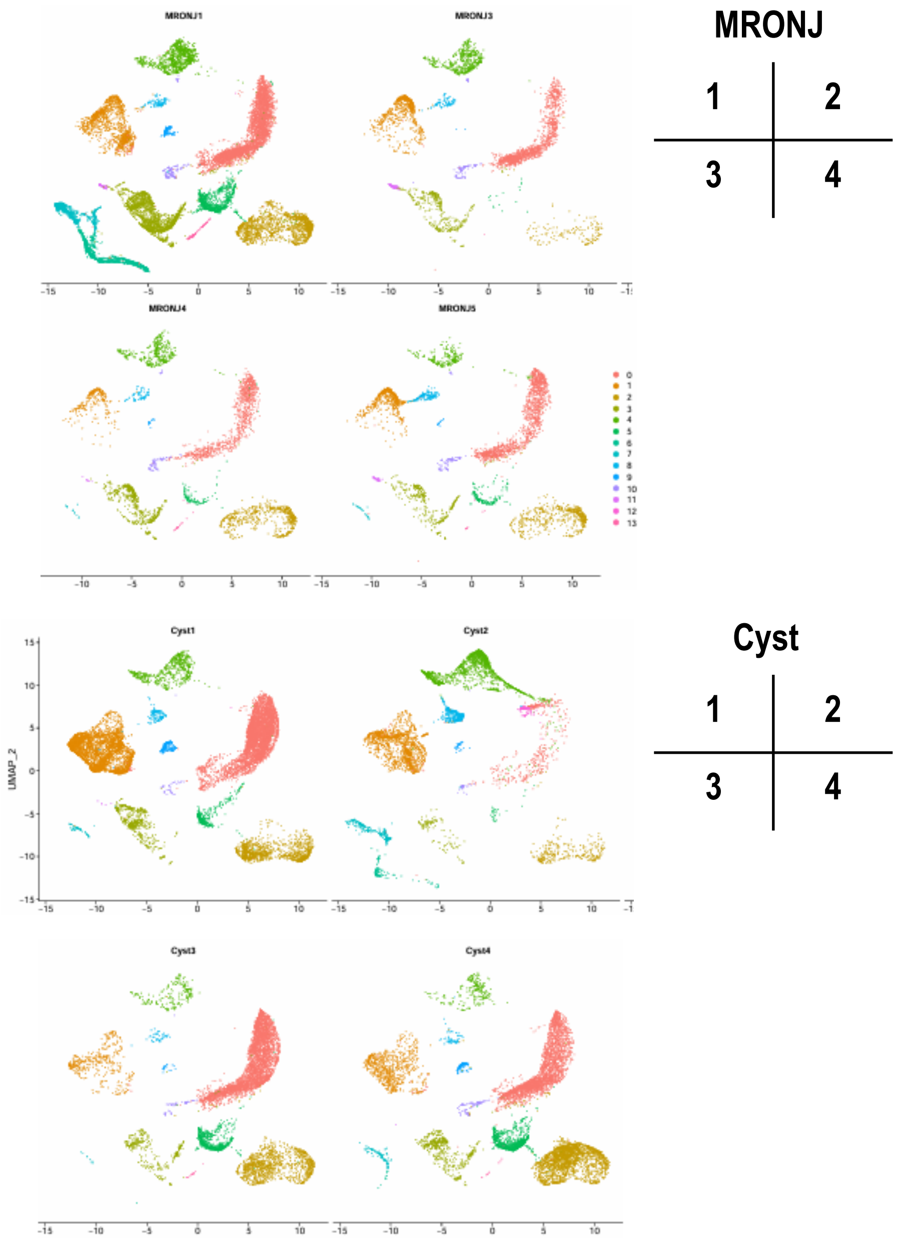

### Supplemental Figure 2

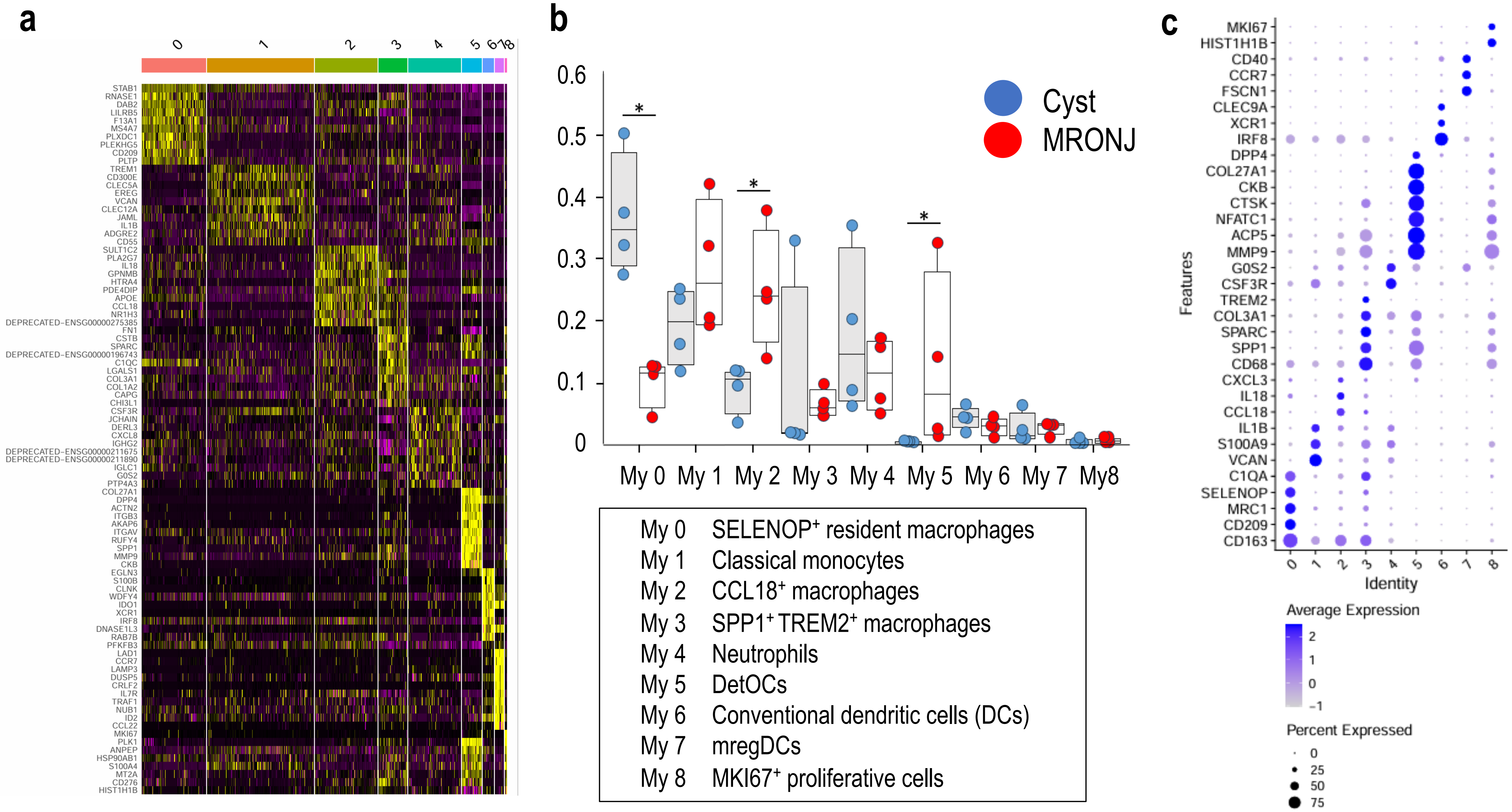

### Supplemental Figure 3

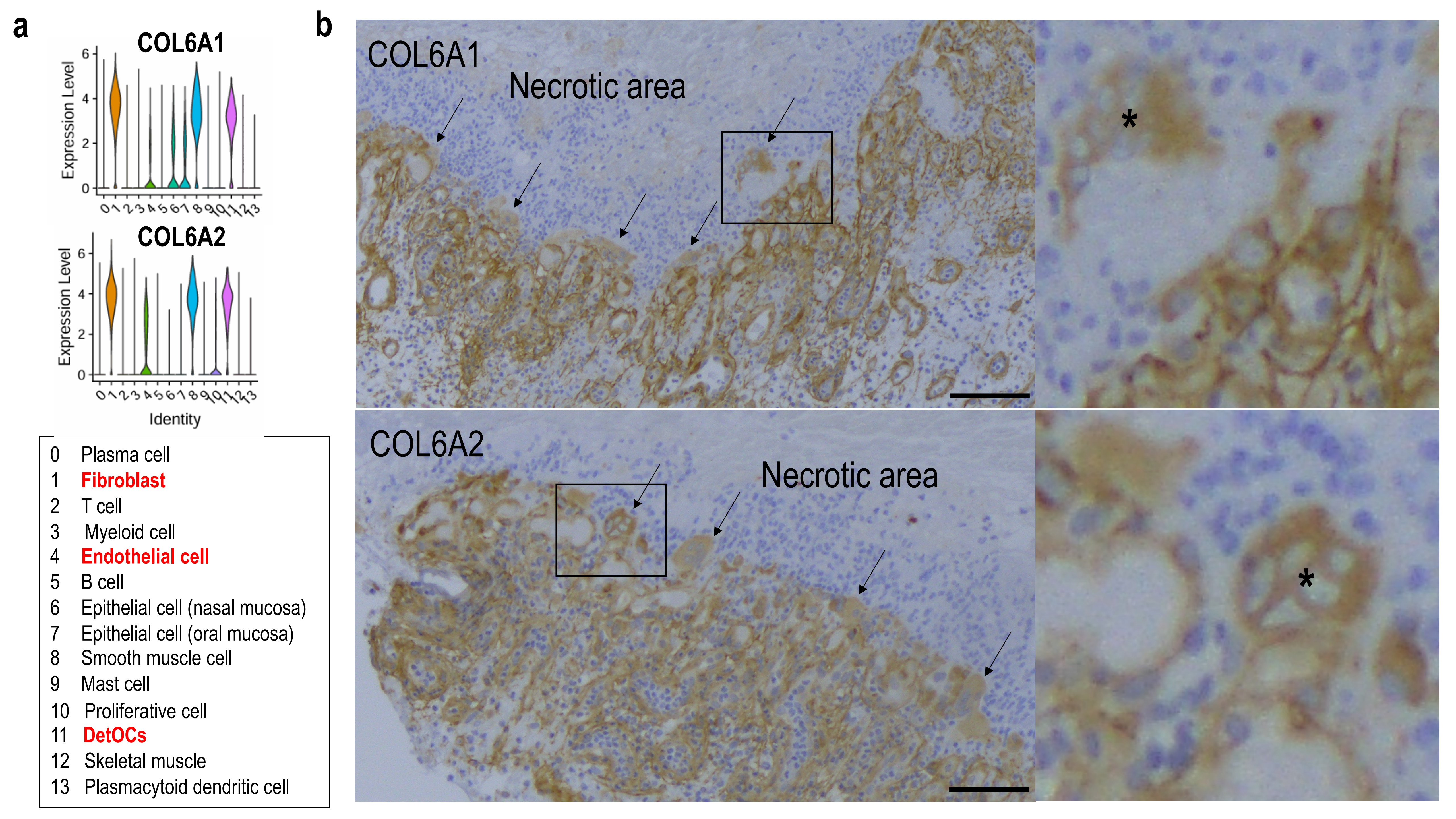

### Supplemental Figure 4

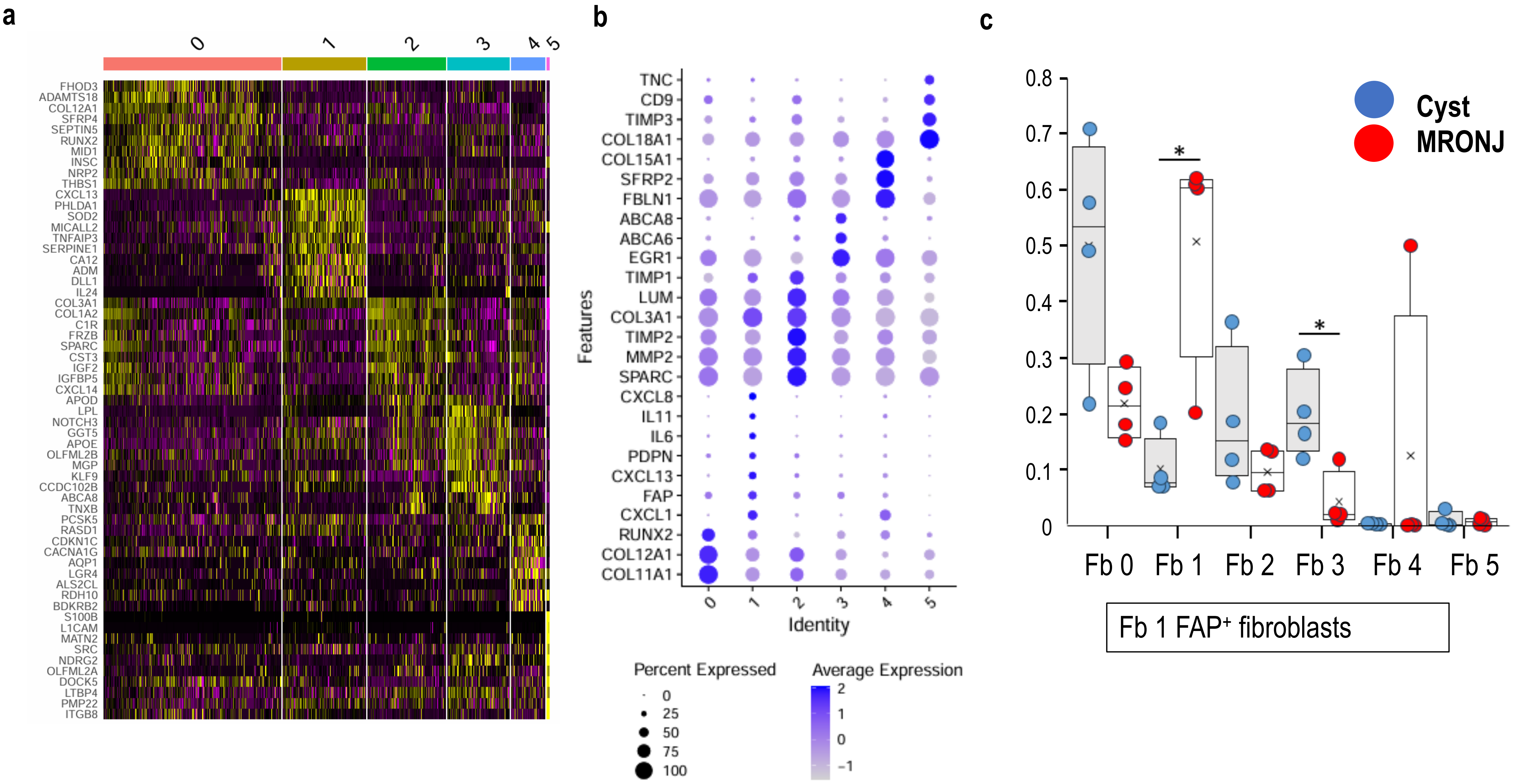
